# The mammalian protein MTCH1 can function as an insertase

**DOI:** 10.1101/2024.12.11.627916

**Authors:** Anna Roza Dimogkioka, Anni Elias, Doron Rapaport

## Abstract

The outer mitochondrial membrane (OMM) hosts a variety of proteins such as import machineries, enzymes, fission/fusion factors, and pore proteins. In *Saccharomyces cerevisiae*, the MIM complex, consisting of Mim1 and Mim2, mediates the insertion of α-helical proteins into the OMM. Until recently, it was unclear which proteins serve this function in higher eukaryotes. Recent studies identified MTCH2 as the insertase of α-helical proteins into the OMM in mammals. MTCH1 is a paralogue of MTCH2 but its general function and contribution to the biogenesis process are not clear. To better characterize MTCH1, we explored whether MTCH1 or MTCH2 could functionally replace Mim1/Mim2 in yeast. Expression of MTCH1 and MTCH2 in yeast cells lacking Mim1, Mim2, or both revealed that MTCH1, but not MTCH2, could compensate the growth defects upon deleting the MIM complex. Furthermore, MTCH1 could restore the biogenesis of MIM substrates, TOM complex stability, and morphology of mitochondria. These findings indicate that MTCH1 by itself has insertase activity and is a functional homologue of the MIM complex, despite the absence of any evolutionary relation between the mammalian and yeast insertases.

## Introduction

Mitochondria play a central role in cellular metabolism, signalling and energy production. Despite possessing their own genome (mtDNA), mitochondria rely on nuclear encoded proteins for their function. In fact, about 1000 proteins in yeast and 1,500 proteins in humans need to be synthesized in the cytosol before being transported to their specific mitochondrial sub-compartments (Schmidt et al., 2010; Wiedemann & Pfanner, 2017). As double-membrane organelles, mitochondria possess a unique architecture. The outer mitochondrial membrane (OMM) serves as a barrier to the cytosol while acting as a communication hub with the rest of the cell. The inner mitochondrial membrane (IMM) forms invaginations called cristae where oxidative phosphorylation occurs (Xian & Liou, 2021).

The OMM is home to a wide array of proteins, including those involved in protein import, enzymatic functions, mitochondrial fission and fusion, and pore formation (Papić et al., 2013). Dysfunction of these OMM proteins has been linked to human diseases like Alzheimer’s, Parkinson’s and mitochondrial encephalopathy (Bhatia et al., 2021; Henrich et al., 2023; Liu et al., 2021).OMM proteins exhibit distinct structural motifs, primarily categorised as either α- helical or β-barrel proteins. While mitochondrial protein import pathways have been well studied, the mechanism by which α-helical proteins are imported into the OMM has only been partially elucidated.

In *Saccharomyces cerevisiae*, the MIM complex, comprising of Mim1 and Mim2 is responsible for the import of certain proteins containing one or more membrane embedded α-helical segments (Becker et al., 2011). Although the exact complex structure and stoichiometry remain unknown, it is speculated that Mim1 exists in the complex in excess to Mim2 (Dimmer et al., 2012; Popov-Čeleketić et al., 2008) . Tom70 has been shown to be involved in the import of some MIM substrates, mainly acting as a receptor or docking site for precursor proteins such as Ugo1, Om14 and the mammalian protein PBR/TSPO (Becker et al., 2011; Papić et al., 2013; Zhou et al., 2022). Initially, the proteins responsible for the membrane insertion of α-helical proteins in other eukaryotes remained unclear.

In 2018, Vitali *et al*. identified pATOM36 as the functional homologue of the MIM complex in *Trypanosoma brucei* (Vitali et al., 2018). Later, in 2022, Guna *et al*. demonstrated that MTCH2 facilitates the import of a subset of α-helical mammalian OMM proteins (Guna et al., 2022). MTCH2 and its understudied paralogue MTCH1 are members of the SLC25 membrane carrier family. Unlike most SLC25 proteins, which localise to the IMM, MTCH1 and MTCH2 are situated in the OMM (Ruprecht & Kunji, 2020). Both proteins carry six putative transmembrane α-helices with MTCH2 having 303 amino acid residues while MTCH1 harbours a total of 389 a.a. residues (Fig. 1A). Most of this difference is due to an extended N-terminal domain of MTCH1, which is predicted by Alpha Fold to be unstructured (Fig. 1B). Of note, MTCH1 was recently identified as a potential anti-ferroptosis factor in cervical cancer (Wang et al., 2023). However, so far, much remained unknown about MTCH1, and although its down-regulation together with reduced levels of MTCH2 resulted in compromised insertion of some OMM proteins (Guna et al., 2022), its single knock-down did not affect biogenesis of OMM proteins (Muthukumar et al., 2024; Wang et al., 2023). MTCH2, on the other hand, has been extensively studied, with suggested roles in lipid homeostasis, mitochondria fusion, and apoptosis ((Labbé et al., 2021; Strauss et al., 2024; Zaltsman et al., 2010)and more recently lipid scrambling (Bartoš et al., 2024; Li et al., 2024). Employing coarse-grained molecular dynamics simulations, Li et al. (2024), concluded that MTCH1 also can have scrambling activity.

**Figure 1.**
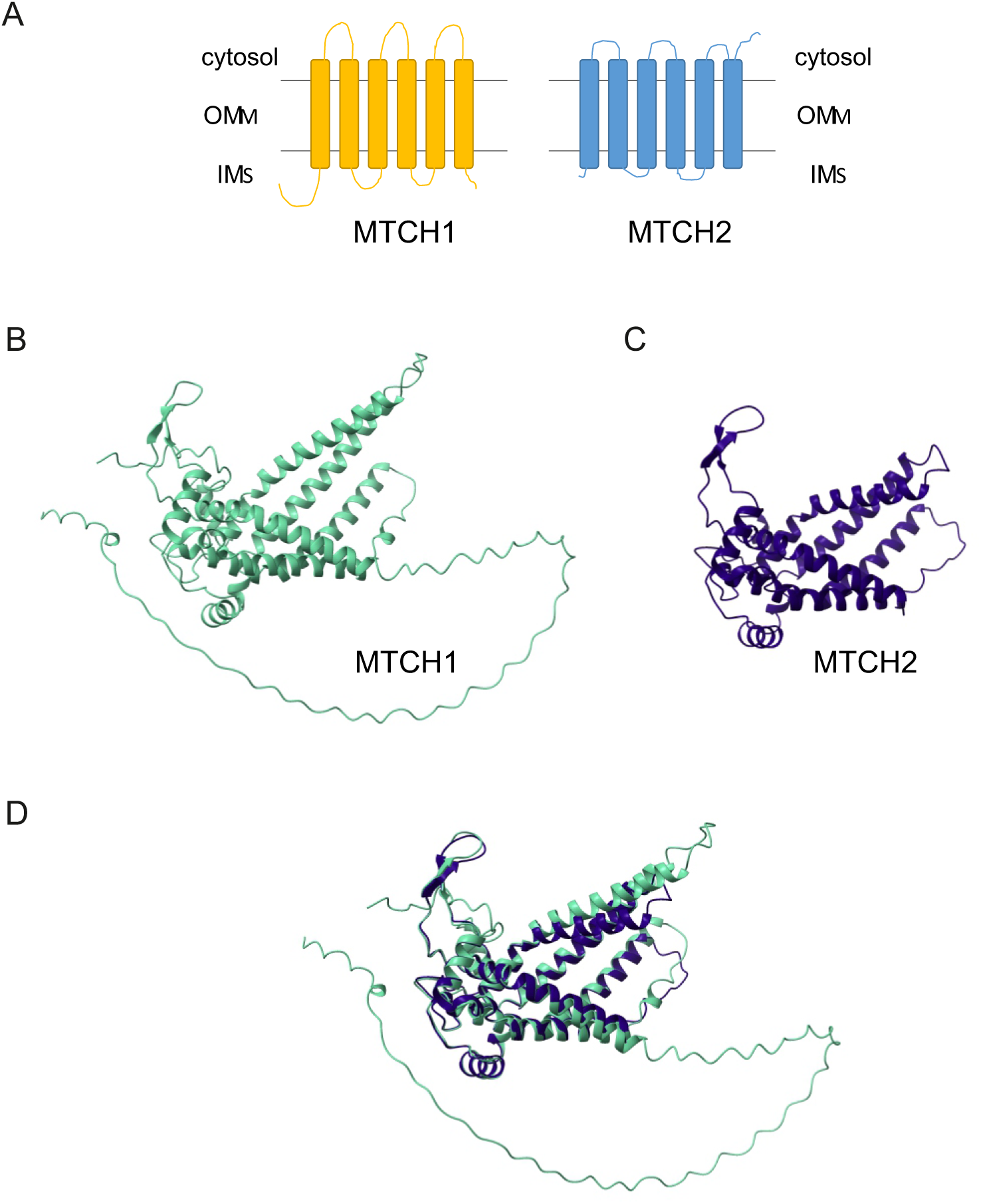
**MTCH1 and MTCH2 are multispan proteins that share structural similarity**. **(A)** Schematic representation of MTCH1 (yellow) and MTCH2 (blue). **(B and C)** Structural prediction of MTCH1 (B, UniProt: Q9NZJ7) and MTCH2 (C, UniProt: Q9Y6C9), adapted from Alpha Fold. **(D)** Overlay of MTCH1 and MTCH2 structural predictions. MTCH1 and MTCH2 structural predictions were aligned using Chimera X.

Notably, MTCH1 and MTCH2 share no sequence similarity with Mim1, Mim2, or pATOM36. While certain SLC25 carrier family proteins are conserved between yeast and mammals, MTCH1 and MTCH2 are not. Given these differences, we sought to investigate whether MTCH1 and/or MTCH2 could functionally complement the absence of Mim1, Mim2, or both in yeast. To that end, we expressed the former proteins in yeast strains mutated for Mim components. Remarkably, MTCH1, but not MTCH2, was able to complement the growth defects associated with the loss of Mim1 and Mim2. Furthermore, MTCH1 restored in the MIM mutated cells MIM substrate biogenesis, TOM complex stability, and mitochondrial morphology. These findings suggest that despite the absence of evolutionary conservation between these mammalian and yeast proteins, MTCH1 possesses inherent insertase activity and functions as a homologue of the MIM complex.

## Results

### MTCH1 Complements the Growth Defects in Cells Lacking Mim1, Mim2 or Both

Given that both MTCH1 and MTCH2 have been implicated in the biogenesis of α-helical OMM proteins (Guna et al., 2022), we sought to investigate their effects when expressed in *S. cerevisiae*. Structurally, both MTCH1 and MTCH2 possess six putative transmembrane α- helices (Fig. 1A); however, MTCH1 is larger due to a long unstructured N-terminal domain (Fig. 1B & C). Structural analysis via AlphaFold predicts tighter α-helices in MTCH2 compared to MTCH1, a difference confirmed by *in silico* structural alignment (Fig. 1D). Since the structure of the MIM complex remains unresolved, we were unable to directly compare it to MTCH1 and MTCH2.

To test whether MTCH1 and/or MTCH2 could functionally substitute the MIM complex *in vivo*, we wished to express these proteins in wild type (WT), or in the single deletion strains *mim1Δ* and *mim2Δ,* as well as the double deletion one *mim1/2ΔΔ*. To that goal, the sequences of MTCH1 and MTCH2 were codon optimised for yeast expression and C-terminally tagged with HA to facilitate detection. As controls, the empty yeast expression plasmid pRS426 and pRS426 plasmid encoding either Mim1 or Mim2 were transformed into the aforementioned strains. Growth of these strains was monitored at either 30⁰C or 37⁰C on solid and liquid media, using either glucose or galactose as carbon sources.

Initial experiments with C-terminally HA-tagged MTCH1 and MTCH2 revealed partial growth complementation of the *mim1Δ*, *mim2Δ* and *mim1/2ΔΔ* strains by MTCH1 (Suppl. Fig. 1). Surprisingly, MTCH2, which had been identified as a key insertase for α-helical OMM proteins (Guna et al., 2022), not only failed to rescue the growth defect but also had a strong inhibitory effect on the growth of all strains, including WT cells (Suppl. Fig. 1).

Considering these unexpected findings, we next wondered whether the HA tag might be responsible for this ‘dominant negative’ phenotype of MTCH2 expression. To test this, we expressed both MTCH1 and MTCH2 without the HA tag. Notably, untagged MTCH1 complemented the absence of Mim1 and Mim2 more effectively than the tagged version. In assays with both solid and liquid media, MTCH1 significantly rescued the growth defects, reaching complementation levels comparable to—but not equal to—those of Mim1 and Mim2 (Fig. 2). Remarkably, MTCH1 alone was able to restore growth in the absence of both Mim proteins, indicating that its activity is independent of preexisting Mim1 or Mim2. Although a very slight negative effect on the growth of WT cells was observed with MTCH1 expression, its expression was beneficial in all other strains (Fig. 2).

**Figure 2.**
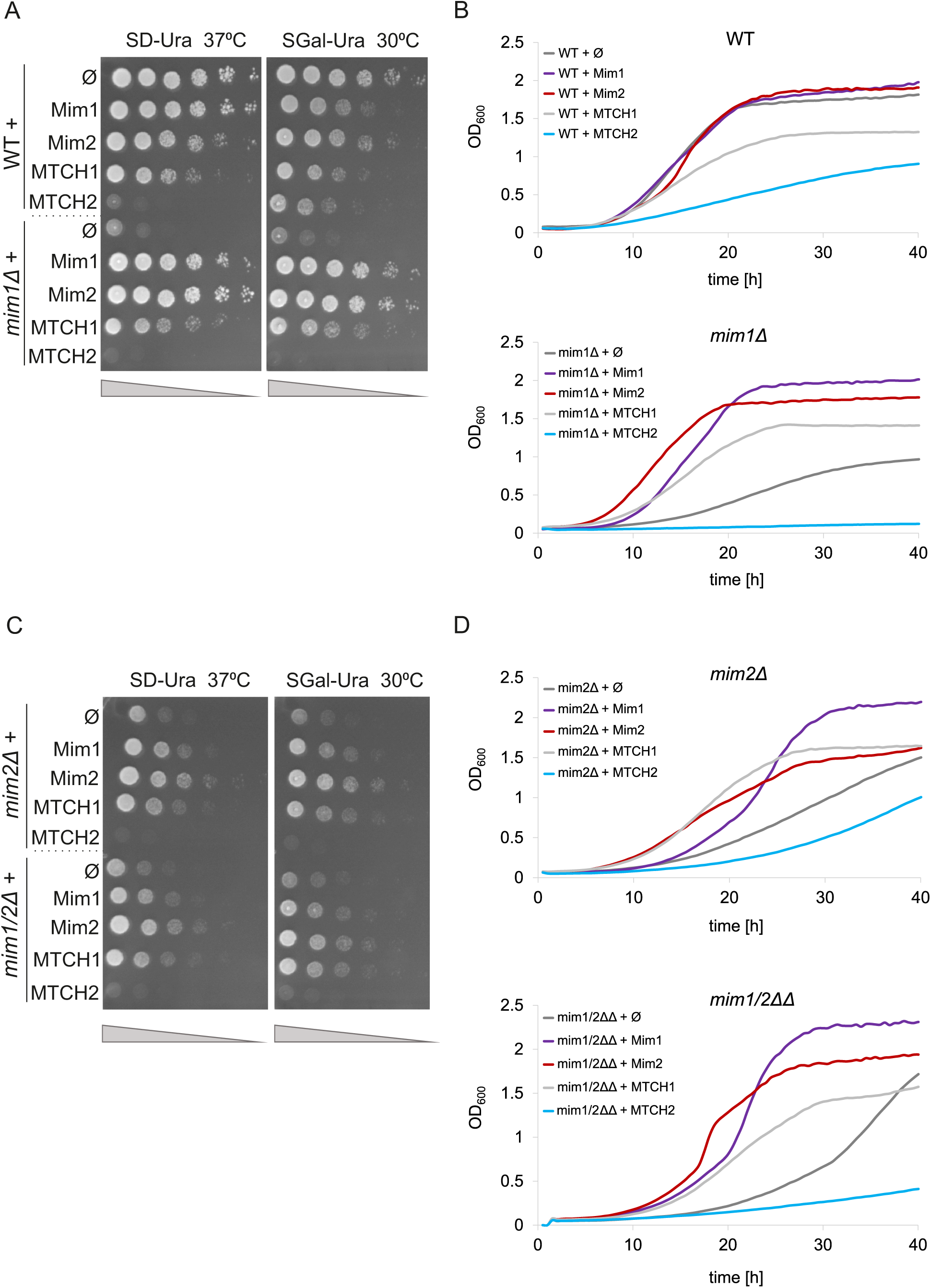
MTCH1 can complement the growth defects in cells lacking Mim1, Mim2 or both. (A and. **C)** The growth of the indicated strains was monitored by drop dilution assay on solid synthetic medium containing either glucose (SD) or galactose (SGal) at either 30⁰C or 37⁰C. The strains were transformed with an empty vector (Φ) or vector encoding the indicated protein. Plates were incubated for 3 days at the indicated temperature before pictures were taken. **(B and D)** The growth of the indicated strains at 37°C in liquid glucose-containing media was observed for 40 hours using OD600 measurements. At the beginning of the measurements (time 0), the strains were diluted to an OD600 of 0.1.

In contrast, MTCH2 exhibited a negative effect on cell growth, even without the HA tag. The most severe growth defect was observed when MTCH2 was expressed in the *mim1Δ* strain (Fig. 2B), with only slight improvements in the *mim2Δ* and *mim1/2ΔΔ* strains (Fig. 2D). In summary, MTCH1 consistently rescued the growth of cells lacking Mim1, Mim2, or both, in both solid and liquid media.

### MTCH1 Restores the biogenesis of MIM Complex Substrate and Reconstitutes TOM Complex Assembly

While MTCH1 successfully complemented the growth defects in strains lacking Mim1 and/or Mim2, it was unclear whether MTCH1 localizes to mitochondria and compensates for other defects associated with the loss of Mim proteins. To confirm mitochondrial localization, we performed subcellular fractionation on cells expressing MTCH1-HA, separating cytosolic, endoplasmic reticulum (ER), and mitochondrial fractions. Hexokinase1 served as a cytosolic marker, Erv2 for the ER, and Hep1 for the mitochondria. As shown in Fig. 3A, MTCH1-HA was found in the mitochondrial fraction alongside Hep1, with small traces in the cytosolic fraction and no detectable presence in the ER fractions, indicating that MTCH1 localizes to yeast mitochondria even in the absence of Mim1 and/or Mim2. Due to the negative effects of MTCH2 on cell growth, isolating mitochondria in MTCH2-expressing cells proved challenging, so further experiments focused solely on MTCH1.

**Figure 3.**
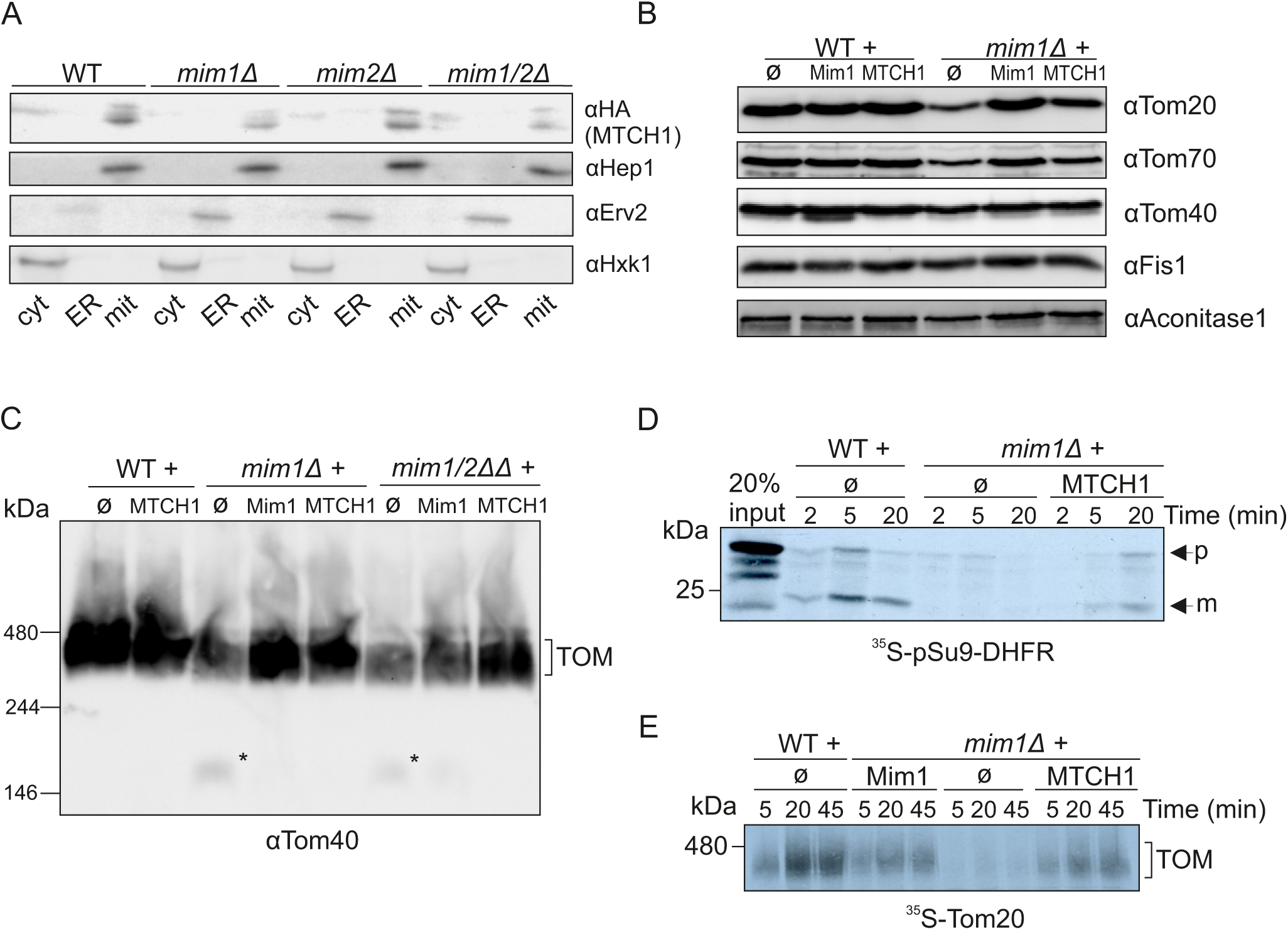
MTCH1 rescues the reduced steady state levels and assembly defects in cells deleted for Mim components. **(A)** Subcellular fractionation of WT, *mim1Δ*, *mim2Δ* and *mim1/2ΔΔ* strains expressing MTCH1-HA. Cytosol, ER and mitochondrial fractions were obtained using differential centrifugation steps. The samples were analysed by SDS-PAGE and immunodecorated with the indicated antibodies; Hep1, a mitochondrial protein; Erv2, an ER protein; Hexokinase (Hxk1), a cytosolic marker. **(B)** Mitochondria (100 µg) isolated from the indicated strains were analysed by SDS-PAGE and immunodecoration with antibodies against the indicated mitochondrial proteins. **(C)** Isolated mitochondria from the indicated strains were solubilised with 1% digitonin. Samples were analysed by BN-PAGE and immunodecorated with antibodies against Tom40. The bands representing the assembled TOM complex are indicated. Asterisk (*) indicates dissociated form of the TOM complex. **(D)** Radiolabelled pSu9-DHFR was synthesised in a cell-free system and imported into mitochondria isolated from the indicated cells. After import for the indicated times, Proteinase K was added. Samples were analysed by SDS-PAGE and autoradiography. **(E)** Radiolabelled Tom20 was imported for the specified time periods into mitochondria isolated from the indicated cells. Samples were analysed via BN-PAGE and autoradiography.

Deletion of Mim1 and Mim2 also leads to reduced levels of several MIM complex substrates, particularly α-helical OMM proteins such as Tom20 and Tom70. We next examined whether MTCH1 could restore the steady-state levels of these α-helical OMM proteins in Mim deletion strains. Additionally, we analysed Tom40, a β-barrel OMM protein, Fis1 that is only very partially requires the MIM complex (Vitali et al., 2020) and Aconitase1, a mitochondrial matrix protein, to ensure that MTCH1’s effects were specific to α-helical OMM proteins. WT and *mim1Δ* deletion strains expressing either empty pRS426 plasmid (Φ) or plasmid-encoded MTCH1 were grown in synthetic media with lactic acid as the carbon source to promote mitochondrial biogenesis. Cells were then lysed, mitochondria were isolated, and their proteins were analysed via SDS-PAGE followed by immunodetection.

The results demonstrated that MTCH1 restored the levels of Tom70 and Tom20 in the deletion strains to levels comparable to those observed in Mim1-expressing cells. Importantly, the expression of MTCH1 in WT cells did not alter the abundance of these proteins (Fig. 3B), indicating that MTCH1 effects are specific to Mim-deficient strains and its activity is not beneficial in the presence of a functional MIM complex.

The loss of Mim1 and Mim2 also disrupts the assembly of the translocase of the outer membrane (TOM) complex, leading to the accumulation of assembly intermediates and reduced levels of the mature TOM complex (Dimmer et al., 2012). This defect arises because several TOM components are α-helical proteins, whose biogenesis is impaired in the absence of the MIM complex. To determine whether MTCH1 expression could correct these assembly defects, we performed blue native (BN)-PAGE analysis on mitochondria isolated from WT and Mim deletion strains transformed with either empty pRS426 plasmid or a plasmid encoding MTCH1. As shown in Fig. 3C, MTCH1 expression alleviated the TOM complex assembly defects observed in *mim1Δ* and *mim1/2ΔΔ* strains. Specifically, MTCH1 reduced the accumulation of TOM complex intermediates (∼100 kDa, indicated by an asterisk) and promoted the formation of the assembled TOM complex (∼440 kDa) to levels comparable to WT mitochondria. These findings indicate that MTCH1 can not only restore the levels of MIM substrates but also improve TOM complex assembly in Mim-deficient strains.

Our findings demonstrate that MTCH1 can rescue growth defects, restore steady-state levels of α-helical mitochondrial outer membrane (MOM) proteins, and reconstitute TOM complex assembly in *mim1Δ* strains. To further investigate the involvement of MTCH1 in the biogenesis of MIM complex substrates, we employed a cell-free translation system to synthesise radiolabelled substrate proteins. These labelled substrates were incubated at 25°C for various time periods with mitochondria isolated from either wild-type (WT) or *mim1Δ* cells.

Although not a specific substrate of the MIM complex, pSu9-DHFR import serves as a general indicator of mitochondrial protein import efficacy, which is known to be (indirectly) compromised when the MIM complex is mutated (Mnaimneh et al., 2004). Thus, we examined the import of pSu9-DHFR, a model fusion protein comprising the presequence of F0-ATPase subunit 9 (pSu9) and mouse dihydrofolate reductase (DHFR). During import, pSu9-DHFR is cleaved by the matrix processing peptidase (MPP) producing a mature form (Fig. 3D and Pfanner et al., 1987). In *mim1Δ* mitochondria, we observed a marked reduction in pSu9-DHFR import, indicating a general defect in mitochondrial protein import. The expression of MTCH1 in *mim1Δ* cells significantly improved pSu9-DHFR import (Fig. 3D).

Finally, we monitored the biogenesis of Tom20, a signal-anchored protein that is embedded in the OMM. Tom20 is known to be a peripheral component of the TOM complex and to heavily relies on the MIM complex for its biogenesis (Hulett et al., 2008; Popov-Čeleketić et al., 2008) Therefore, to monitor the import of newly synthesized radiolabeled Tom20 molecules, Tom20 association with the TOM complex was assessed using BN-PAGE. To account for the weak association of Tom20 with the rest of the TOM complex, gentle solubilization of mitochondria was applied. This BN-PAGE analysis revealed that Tom20 assembly with the TOM complex was impaired in *mim1Δ* mitochondria, with slower import kinetics compared to control organelles. Importantly, MTCH1 expression in *mim1Δ* cells improved the rate of Tom20 integration into the TOM complex similar to the effect observed upon re-introducing Mim1 (Fig. 3E).

### MTCH1 Improves Mitochondrial Morphology in Strains Lacking Mim1

To further confirm that MTCH1 functions as a homolog of the MIM complex, we examined the overall mitochondrial network morphology. Cells lacking Mim1, Mim2, or both exhibit characteristic mitochondrial defects, including fragmentation and punctate structures, in contrast to the tubular mitochondrial network seen in WT cells (Dimmer et al., 2012). To visualize the mitochondrial network, we expressed a fusion protein consisting of green fluorescent protein (GFP) linked to mitochondrial targeting presequence, pSu9 (mt-GFP). We expressed pSu9-GFP in *mim1Δ* cells harbouring an empty pRS426 plasmid, or a plasmid encoding either Mim1 or MTCH1.

As expected, *mim1Δ* cells predominantly displayed a fragmented mitochondrial network with the formation of punctate structures (Fig. 4A). When Mim1 expression was restored, the cells regained a tubular mitochondrial network consistent with the morphology observed in healthy mitochondria (Fig. 4A & B). Strikingly, cells expressing MTCH1 also exhibited a largely restored tubular mitochondrial network, and only approximately 20% of the cells still displayed abnormal mitochondrial morphology (Fig. 4A and B). This major restoration suggests that MTCH1 can support also the recovery of mitochondrial structure.

**Figure 4.**
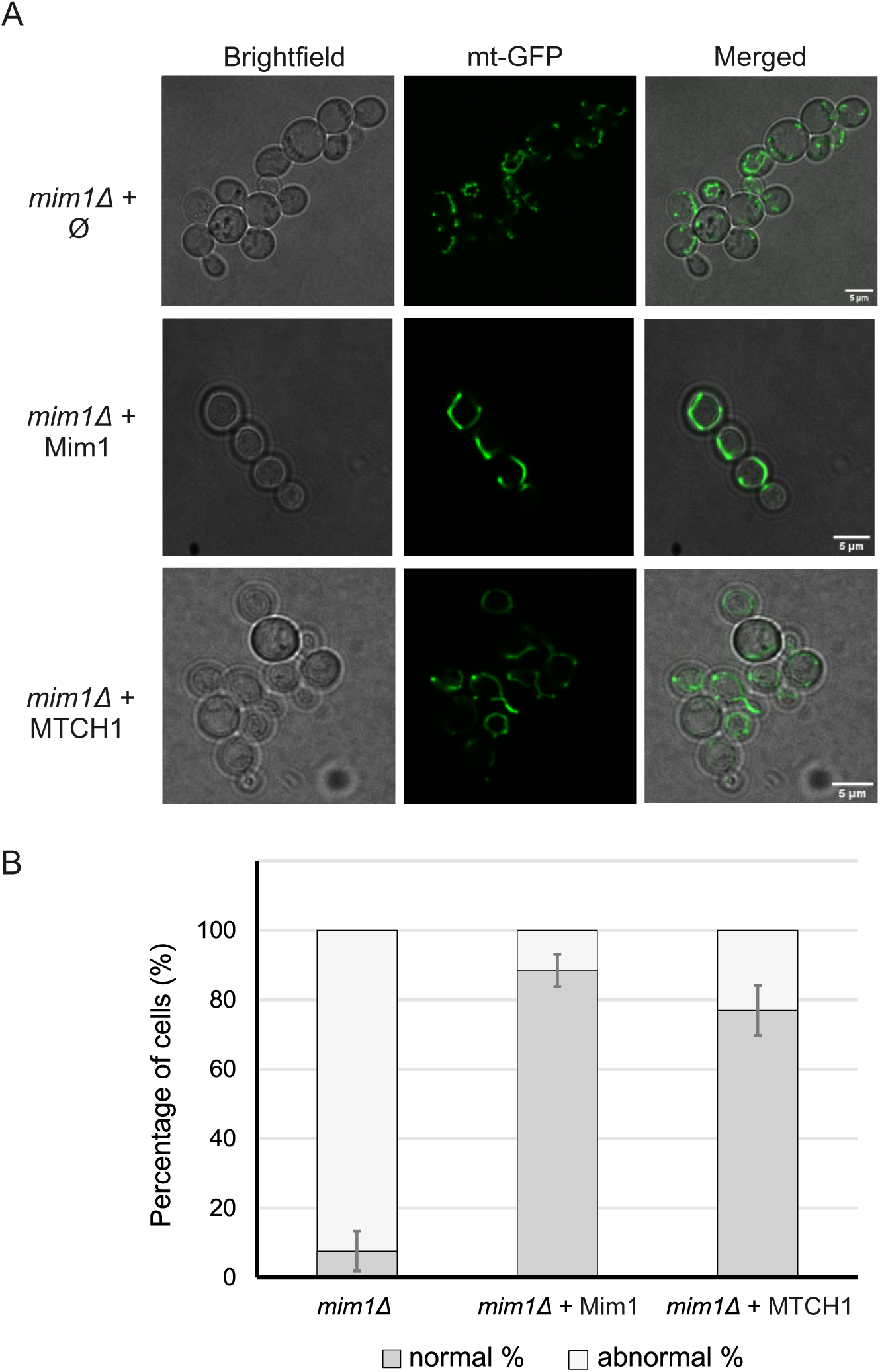
MTCH1 can complement the mitochondrial morphology defect of *mim1Δ* cells. **(A)** *mim1Δ* cells harbouring mitochondria-targeted GFP (mtGFP) were transformed with either an empty plasmid (Ø) as a control (upper panels) or a plasmid encoding either MIM1 or MTCH1 (middle and lower panels, respectively). Cells were analysed by fluorescence microscopy and representative images of the predominant morphology for each strain are shown. Scale bar, 5 µm. **(B)** Statistical analysis of the cells described in (A) where cells with either normal or abnormal mitochondrial morphology were counted. Average values with standard deviation bars of three independent experiments with at least n = 100 cells in each experiment are shown.

## Discussion

MTCH1 has previously been suggested to be involved in the import of α-helical OMM proteins in mammalian cells; however, clear evidence for such a function was missing. Our study demonstrates that MTCH1 alone can functionally complement the absence of Mim1, Mim2, or both in *S. cerevisiae* and fully rescue the import defects in mitochondria lacking a functional MIM insertase. Although MTCH1 and MTCH2 share structural similarities and are members of the same protein family, their effects in yeast are markedly different. Specifically, in contrast to the positive effects of MTCH1, MTCH2 exhibited an inhibitory effect on the growth of WT cells as well as strains lacking Mim proteins.

We can only speculate on the mechanism underling MTCH2’s negative impact on mitochondrial function in yeast. MTCH2 is known to play a key role in apoptosis by recruiting truncated tBID to the OMM, thereby facilitating BAX activation, which leads to OMM permeabilization and cytochrome c release (Raemy et al., 2016; Zaltsman et al., 2010). However, *S. cerevisiae* lacks a direct homolog of tBID or other Bcl-2 family proteins commonly found in mammalian cells (Polčic & Mentel, 2020), even though yeast undergoes regulated cell death. Given that yeast proteomes do not include the apoptotic machinery found in mammals, it is plausible that MTCH2 may retain some apoptotic function in yeast by interacting with yet unidentified, yeast-specific apoptotic factors. Alternatively, MTCH2 may be interfering with the import machinery on the OMM, thereby disrupting mitochondrial homeostasis and leading to impaired growth. Another possibility is that MTCH2 mislocalises to other organelles, such as the ER or peroxisomes, where it could interfere with their function. Obviously, these three potential explanations are not mutually exclusive. Given the complex role of MTCH2 in mitochondrial regulation in mammalian cells, it is very hard to pinpoint the exact reason for its negative effect on yeast cells.

The ability of MTCH1 to functionally replace the MIM complex in yeast mitochondria suggests that MTCH1 possesses distinct properties that make it more adaptable to yeast systems. One possible explanation lies in the structural differences between MTCH1 and MTCH2. MTCH1 contains a large unstructured N-terminal domain that is absent in MTCH2. This region may be crucial for substrate recognition or docking, enabling MTCH1 to act as an efficient insertase in yeast. Additionally, structural prediction analysis has revealed that the α-helices of MTCH2 are more tightly packed compared to those of MTCH1, potentially suggesting that MTCH1’s conformation may be better suited for integration into the yeast mitochondrial membrane.

Moreover, while structural data for the MIM complex remain unavailable, future studies comparing the structure of MTCH1 to that of the MIM complex could reveal some evolutionary conservation. It is possible that MTCH1 has more structural similarity to the MIM complex than MTCH2, allowing it to perform the same roles in protein import. MTCH2, on the other hand, may have evolved later to fulfil more specialized or complementary functions in complex tissues and cells of mammalian organisms. The lack of sequence and apparent structural similarity between Mim subunits and MTCH1 despite their related functions is in line with the previous identification of the protein pATOM36 as the functional equivalent of the MIM complex in *T. brucei* (Vitali et al., 2018). Also, pATOM36 and the Mim components do not share any sequence and structural resemblance. Hence, it seems that the different insertases of the OMM are the result of convergent evolution.

Our novel findings open avenues for further exploration into the evolutionary significance of functional conservation in mitochondrial biogenesis. To provide deeper insights into the functional redundancy and evolutionary history of mitochondrial insertases across species, future studies might test the reciprocal complementation, where both components of the MIM complex (Mim1/2) are co-expressed in human cell lines lacking MTCH1 and/or MTCH2. It is hard to predict whether such co-expression will restore function in a manner similar to our findings in yeast.

In summary, our current findings provide clear evidence for the capacity of the mammalian protein MTCH1 to act as an insertase for mitochondrial outer membrane proteins.

## Materials and Methods

### Yeast strains and growth conditions

Yeast strains of the genetic background W303α that were described before were used in this study (Dimmer et al., 2012). Yeast cells were grown at either 30⁰C or 37⁰C in synthetic medium S-uracil (1.9 g yeast nitrogen base without ammonium sulphate, 5 g ammonium sulphate, 55 mg adenine sulphate) supplemented with 100x stock of amino acids and glucose (2%) or galactose (2%). For mitochondrial isolation, cells were grown in S-uracil media supplemented with lactate (2%). Yeast strains were transformed with the desired plasmid(s) using the lithium acetate method.

### Recombinant DNA methods

For better expression in yeast cells, the DNA encoding sequences of both MTCH1 and MTCH2 were optimised for yeast codon usage utilizing the Eurofins Genomics codon optimisation tool. Both genes were cloned into the pRS426 and pYX142 vectors for yeast expression purposes with EcoRI and BamHI as the restriction sites. A detailed lists of primers and plasmids used in this study are included in Tables S1 and S2, respectively.

### Cell growth assays

For growth assay on solid medium (drop dilution assays), yeast strains were grown in SD-ura medium to an OD600 = 1. The cells were then serially diluted five times in fivefold increment. Next, 5 µl from each dilution was placed onto the solid medium and the plates were incubated at 30⁰C or 37⁰C for several days. For the liquid growth curve assays, yeast strains were grown to an OD600 = 1 and diluted to OD600 = 0.1. Triplicates of each strain were loaded onto a 96- well Microtest plate (SARSTEDT) and the OD600 was measured every 30 min for up to 60 hours using the SPECTROstar Nano plate reader. Analysis of the growth curves was performed using the SPECTROstar Nano MARS software.

### Biochemical methods

#### Isolation of Mitochondria

Yeast cells were grown in liquid media (volume of 2-6 L) to logarithmic phase. The cells were harvested (3000xg, 5 min, RT), resuspended in DTT buffer and incubated at 30 °C for 15 min. Cells were harvested (2000xg, 5 min, RT), washed once with spheroplasting buffer (1.2 M Sorbitol, 20 mM KPI, pH 7.2), harvested again and resuspended in spheroplasting buffer with Zymolyase (5 mg/g of cells) and incubated at 30°C for 1 hour.

Further steps were carried out on ice. Spheroplasts were homogenized in homogenization buffer (0.6 M Sorbitol, 10 mm Tris, pH 7.4, 1 mM EDTA, 0.2% fatty acid-free BSA with 2 mM phenylmethylsulfonyl fluoride (PMSF) using a dounce homogenizer to obtain a cell lysate. Cell debris and nuclei were removed by two clarifying spins (2000 x g, 10 min, 4°C). The supernatant (cytosol + organelles) was centrifuged (18,000xg, 15 min, 4°C) to pellet crude mitochondria. The resulting post nuclear supernatant (PNS) consisted of ER/microsomal and cytosolic fractions. The crude mitochondria were washed twice with SEM buffer (250 mM Sucrose, 1 mM EDTA, 10 mM MOPS) containing 2 mM PMSF and were pelleted again (18,000xg, 15 min, 4°C).

#### Subcellular fractionation

All the steps were carried out at 4°C. Whole cell lysate and crude mitochondria were obtained as described above. To further purify mitochondria from potential contaminants, the mitochondrial fraction was layered on a Percoll gradient (25% Percoll, 2 M sucrose, 100 mM MOPS/KOH pH 7.2, 100 mM EDTA, 200 mM PMSF) and centrifuged (80,000 x g, 45 min, 4°C). Highly pure mitochondria were found as a brownish layer close to the bottom of the tube and was removed carefully with a Pasteur pipette. The mitochondria were washed several times with SEM buffer containing 2 mM PMSF and pelleted again (18,000xg, 15 min, 4°C).

To isolate ER/microsomal and cytosolic fractions, 20 ml of PNS were clarified (18,000 x g, 15 min, 4°C) and centrifuged (200,000xg, 1 hour, 4°C). The supernatant contained the cytosolic fraction. The brownish sticky pellet (consisting of ER) was resuspended in 2 ml of SEM buffer containing 2 mM PMSF and homogenized with a dounce homogenizer. The sample was centrifuged (18,000xg, 20 min, 4 C) to obtain ER/microsomes in the supernatant.

The obtained fractions were precipitated with chloroform-methanol mixture and the pellet was resuspended in 2x sample buffer (125 mM Tris pH 6.8, 4% SDS, 20% glycerol, 10% β-ME, 2 mg/mL bromophenol blue) to obtain protein concentration of 2 mg/ml. Samples were heated at 95°C for 10 min and further analysed by SDS-PAGE and immunoblotting. Table S4 indicates the antibodies used in the current study.

#### Protein analysis

SDS-PAGE was used to separate protein samples. Samples were resuspended in 2x Laemmli buffer with 5% 2-Mercaptoethanol, boiled at 95⁰C for 10 min and loaded for analysis onto gels ranging from 10-15% acrylamide. Proteins were subsequently transferred onto nitrocellulose membranes and immunodecorated. Table S3 includes a list of antibodies used in this study. Quantification of western blots was performed using the AIDA software.

To analyse proteins in their native state, BN-PAGE was used; isolated mitochondria were resuspended in 1% digitonin (digitonin to protein ratio of 6:1) in SEM buffer. After 30 min incubation, the samples were centrifuged (30,000xg, 20 min, 2⁰C) and the supernatant was mixed with 10x BN loading dye. After the separation on the gel, proteins were blotted onto polyvinylidene difluoride (PVDF) membrane and immunodetection proceeded as for SDS-PAGE.

#### In vitro import assay

For the *in vitro* import assay with radiolabelled proteins and isolated mitochondria, a previously published protocol was used (Mokranjac & Neupert, 2007). In this study, a mixture of radiolabelled cysteine and methionine was used during the translation of the proteins. Accordingly, excess of unlabelled methionine and cysteine were used at the end of the translation step. The import of pSu9-DHFR was followed by Proteinase K (PK, 50 µg/ml) treatment and the activity of PK was inhibited using 5 mM PMSF. The import of Tom20-3xHA was followed by BN-PAGE. After import at 25⁰C for various time periods, the isolated mitochondria were collected by centrifugation (13,200xg, 10 min, 2⁰C) and then resuspended in SEM buffer with 1% digitonin. Further analysis was as described above.

#### Microscopy

To study mitochondrial morphology of yeast cells via fluorescent microscopy, yeast strains were transformed with the pYX122-pSu9-GFP plasmid resulting in GFP stained mitochondria. Cells were grown to an OD600 of 0.8-1.0, harvested and mixed with 1% (w/v) low melting agarose to limit their movement on the coverslip. Fluorescent images were taken using a confocal spinning disk microscope Zeiss Axio Examiner Z1 with a CSU-X1 real-time confocal system (Visitron) and SPOT Flex charge-coupled device camera. Fluorescent images were analysed using ImageJ (Fiji).

## Acknowledgments

We thank E. Kracker for excellent technical assistance and K.S. Dimmer for helpful discussions. This work was supported by the Deutsche Forschungsgemeinschaft (RA 1028/11- 1 to D.R. and through Research Training Group 2364/2 to A.R.D. and D.R.),

The authors declare no competing financial interests.

## Author contributions

A.R.D. designed and conducted experiments, A.E. performed experiments; D.R. designed experiments and analysed data, A.R.D. and D. R. wrote the initial version of the manuscript. All authors read and contributed to the final manuscript.

## Supplementary Tables

**Suppl. Figure 1.**
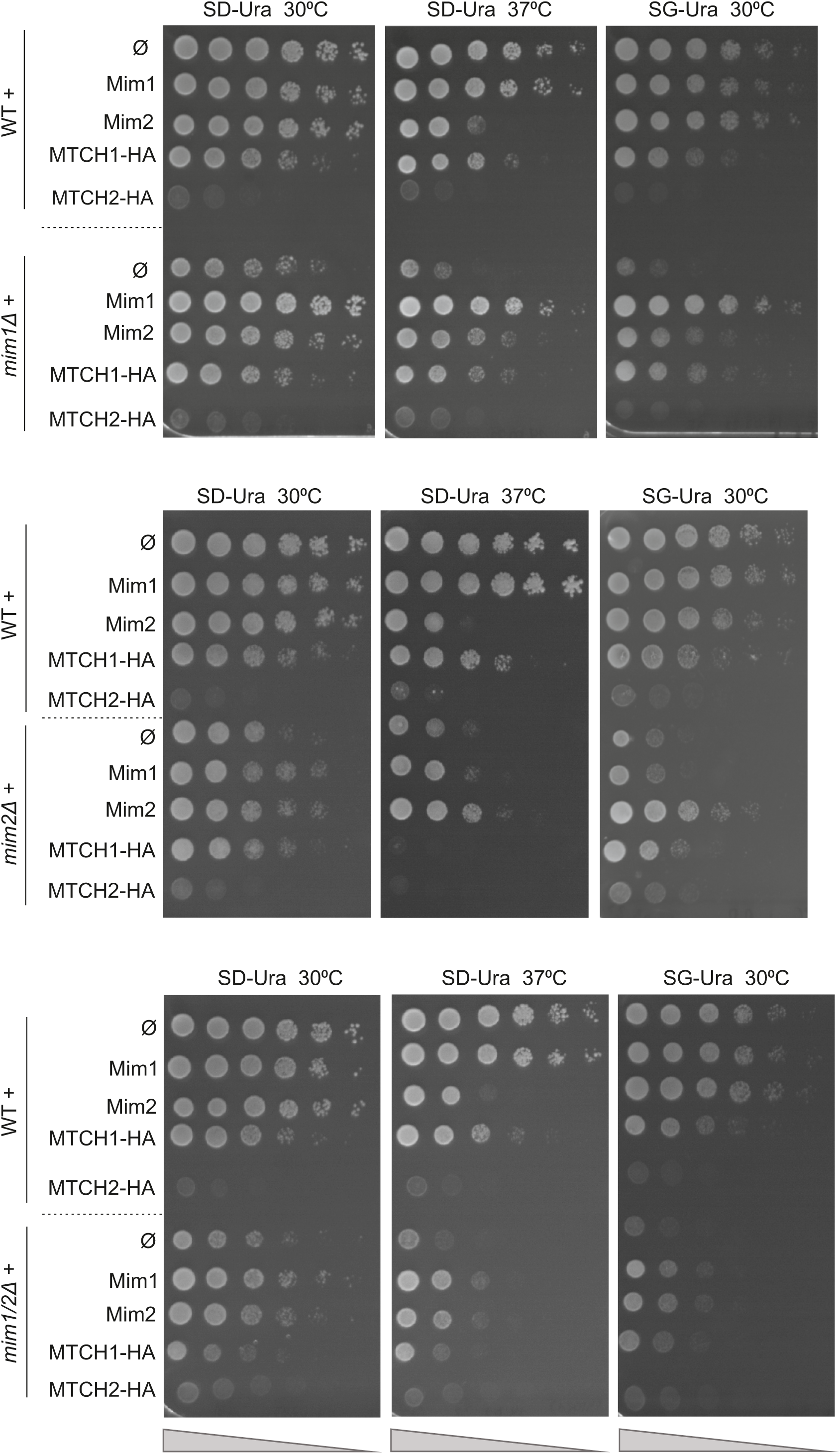
HA-tagged MTCH1 can rescue growth retardation in cells mutated for Mim components. The growth of the indicated strains was monitored by drop dilution assay on solid synthetic medium containing either glucose (SD) or glycerol (SG) at either 30⁰C or 37⁰C. The strains were transformed with an empty vector (Φ) or vector encoding the indicated protein. Plates were incubated for 3 days at the indicated temperature before pictures were taken.

**Table S1:**
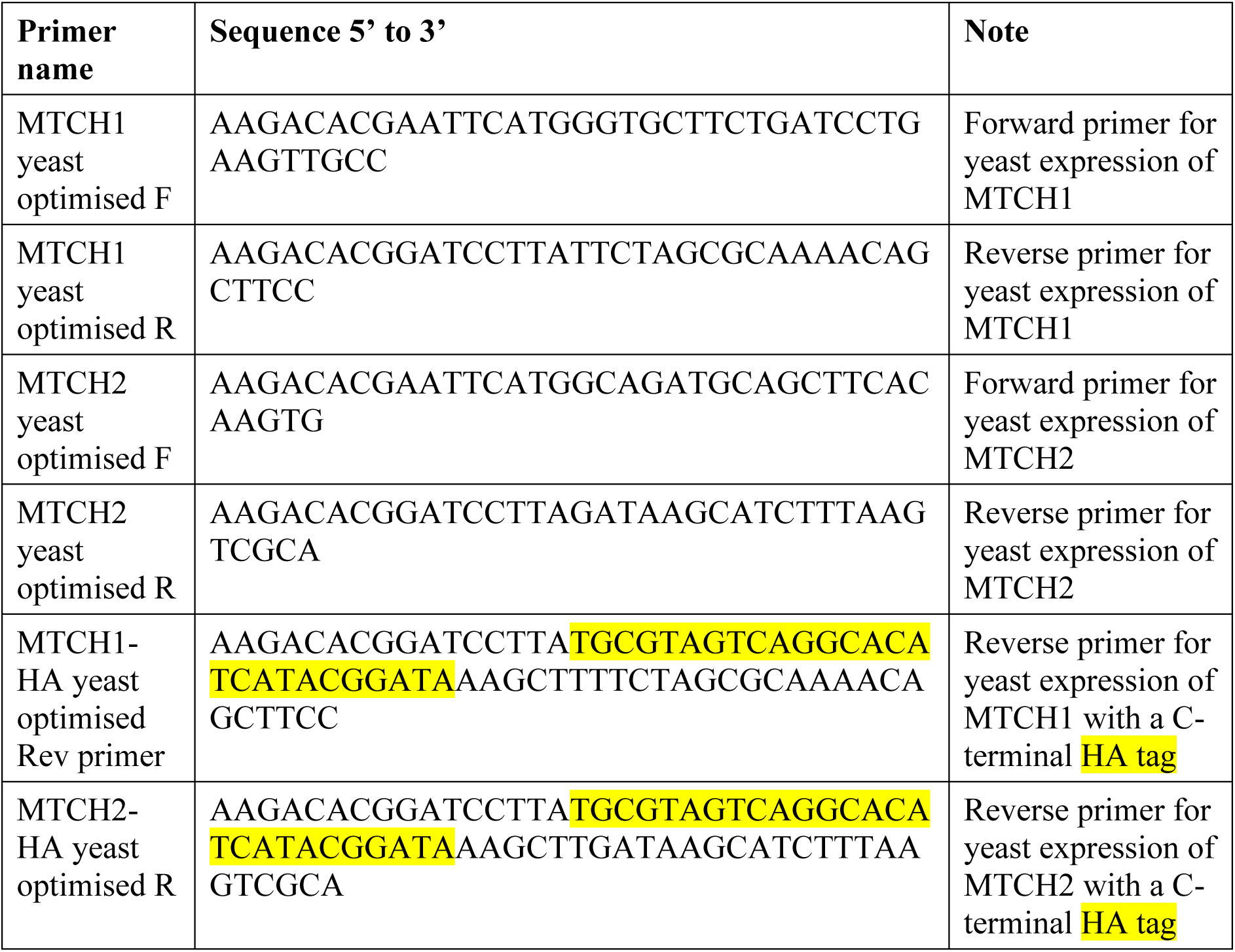
Primers used in this study.

**Table S2:** Plasmids used in this study.

| **Plasmid** | **Promoter** | **Markers** | **Purpose** | **Source** |
| --- | --- | --- | --- | --- |
| pRS426 | TPI | Amp<sup>R</sup>, URA3 | Yeast expression | Lab Stock |
| pRS426 Mim1 | TPI | Amp<sup>R</sup>, URA3 | Yeast expression | This study |
| pRS426 Mim2 | TPI | Amp<sup>R</sup>, URA3 | Yeast expression | This study |
| pRS426 MTCH1 | TPI | Amp<sup>R</sup>, URA3 | Yeast expression | This study |
| pRS426 MTCH2 | TPI | Amp<sup>R</sup>, URA3 | Yeast expression | This study |
| pRS426 MTCH1 | TPI | Amp<sup>R</sup>, URA3 | Yeast expression | This study |
| pRS426 MTCH2 | TPI | Amp<sup>R</sup>, URA3 | Yeast expression | This study |
| pYX142 | TPI | Amp<sup>R</sup>, LEU2 | Yeast expression | Lab Stock |
| pYX142 MTCH1 | TPI | Amp<sup>R</sup>, LEU2 | Yeast expression | This study |
| pYX142 MTCH2 | TPI | Amp<sup>R</sup>, LEU2 | Yeast expression | This study |
| PYX122 pSu9-GFP | TPI | Amp<sup>R</sup>, HIS3 | Yeast expression | Lab stock |
| pGEM4 pSu9-DHFR | SP6/T7 | Amp<sup>R</sup> | In vitro transcription | Lab Stock |
| pGEM4 Om14 | SP6/T7 | Amp<sup>R</sup> | In vitro transcription | Lab Stock |
| pGEM4 3xHA-Tom20 | SP6/T7 | Amp<sup>R</sup> | In vitro transcription | Lab Stock |
| pSP6 PBR | SP6 | Amp<sup>R</sup> | In vitro transcription | Lab Stock |

**Table S3:** Antibodies used in this study.

| **Antibody** | **Source or reference** | **Dilution** |
| --- | --- | --- |
| Polyclonal rabbit αFis1 | Lab Stock | 1:1000 |
| Polyclonal rabbit αTom20 | Lab Stock | 1:1000 |
| Polyclonal rabbit αTom40 | Lab Stock | 1:1000 |
| Polyclonal rabbit αTom70 | Lab Stock | 1:500 |
| Polyclonal rabbit αAcconitase1 | Lab Stock | 1:1000 |
| Polyclonal rabbit αErv2 | Lab of Roland Lill | 1:1000 |
| Polyclonal rabbit αHep1 | Lab Stock | 1:1000 |
| Polyclonal rabbit αHexokinase1 | Bio-Trend #100-4159 | 1:1000 |
| Polyclonal rat anti-HA | Roche #11867423001 | 1:1000 |
| Goat anti-rabbit IgG HRP conjugate | BioRad #1721019 | 1:5000 |

## Notes

### Competing Interest Statement

The authors have declared no competing interest.

